# Potential to Enhance Large Scale Molecular Assessments of Skin Photoaging through Virtual Inference of Spatial Transcriptomics from Routine Staining

**DOI:** 10.1101/2023.07.30.551188

**Authors:** Gokul Srinivasan, Matthew Davis, Matthew LeBoeuf, Michael Fatemi, Zarif Azher, Yunrui Lu, Alos Diallo, Marietta Saldias Montivero, Fred Kolling, Laurent Perrard, Lucas Salas, Brock Christensen, Scott Palisoul, Gregory Tsongalis, Louis Vaickus, Sarah Preum, Joshua Levy

## Abstract

The advent of spatial transcriptomics technologies has heralded a renaissance in research to advance our understanding of the spatial cellular and transcriptional heterogeneity within tissues. Spatial transcriptomics allows investigation of the interplay between cells, molecular pathways and the surrounding tissue architecture and can help elucidate developmental trajectories, disease pathogenesis, and various niches in the tumor microenvironment. Photoaging is the histological and molecular skin damage resulting from chronic/acute sun exposure and is a major risk factor for skin cancer. Spatial transcriptomics technologies hold promise for improving the reliability of evaluating photoaging and developing new therapeutics. Current challenges, including limited focus on dermal elastosis variations and reliance on self-reported measures, can introduce subjectivity and inconsistency. Spatial transcriptomics offer an opportunity to assess photoaging objectively and reproducibly in studies of carcinogenesis and discern the effectiveness of therapies that intervene on photoaging and prevent cancer. Evaluation of distinct histological architectures using highly-multiplexed spatial technologies can identify specific cell lineages that have been understudied due to their location beyond the depth of UV penetration. However, the cost and inter-patient variability using state-of-the-art assays such as the 10x Genomics Spatial Transcriptomics assays limits the scope and scale of large-scale molecular epidemiologic studies. Here, we investigate the inference of spatial transcriptomics information from routine hematoxylin and eosin-stained (H&E) tissue slides. We employed the Visium CytAssist spatial transcriptomics assay to analyze over 18,000 genes at a 50-micron resolution for four patients from a cohort of 261 skin specimens collected adjacent to surgical resection sites for basal and squamous keratinocyte tumors. The spatial transcriptomics data was co-registered with 40x resolution whole slide imaging (WSI) information. We developed machine learning models that achieved a macro-averaged median AUC and F1 score of 0.80 and 0.61 and Spearman coefficient of 0.60 in inferring transcriptomic profiles across the slides, and accurately captured biological pathways across various tissue architectures.

## 1. Introduction

Spatial transcriptomics is an innovative and rapidly evolving field in biomedical research that combines the power of genomics and spatial mapping techniques to gain insights into the spatial organization of gene expression within complex tissues, such as the skin. By providing a detailed view of gene expression patterns in relation to cellular and tissue architecture, spatial transcriptomics has quickly become a valuable tool for biomedical research, including dermatological research.

The skin is the largest organ in the body, composed of multiple cell types that each play a crucial role in maintaining its structure and function. Though traditional genomic analysis techniques, such as bulk-RNA sequencing (RNA-seq), and disaggregated techniques such as single cell RNA sequencing (scRNA-seq), have provided valuable information about cellular heterogeneity and disease progression, they lack the ability to assess localized gene expression patterns that may relate with cell-cell interactions and architecture to support tissue function. Spatial transcriptomics approaches uniquely allow researchers to examine gene expression patterns within their anatomical and histological context, enabling a deeper understanding of the underlying molecular mechanisms driving skin biology, carcinogenesis, and disease progression.

An important potential application of spatial transcriptomics in dermatology is to advance emerging study of skin aging.^1^ The skin serves as a barrier between the environment and the body where it is exposed to near-constant insults including ultraviolet radiation (UVR), mechanical stress, and toxicants.^2^ This exposure, along with genetic influences, combine to induce skin damage, reduced function, and, ultimately, a characteristic wrinkled appearance of the skin largely reflecting degradation of the collagen matrix.^3^ More recently, Zou et al. created a single-cell transcriptomic atlas of human skin aging using eyelid tissue and identified cell-type-specific associations with human skin aging.^1^ Further characterization of cellular changes that incorporate spatial information in skin can inform therapeutic strategies and interventions to combat age-related skin alterations and disease.

Currently, spatial transcriptomics technologies at whole transcriptomic-level multiplexing is incredibly costly and prone to several sources of variation (e.g., within/between-subject variation), limiting broad application. Recently, deep learning models have been proposed as a cost-saving alternative to predict spatial gene expression from routine tissue stains.^4–6^ For instance, the DeepSpaCE approach includes convolutional neural networks (CNNs) for spatial gene cluster and gene expression prediction in human breast cancer tissue sections.^4^ Another modeling paradigm aimed at predicting spatial gene expression across breast tumor and cutaneous tumor data used a mix of transformer and graph neural network-based approaches.^6^ In addition, the performance of several different modeling approaches for spatial gene expression prediction in tissue was recently compared using stage-III (pT3) colorectal tumors. Though these studies demonstrate the potential to infer spatial expression patterns using histomorphological data, several crucial questions remained unanswered, including the applicability of these methods to non-cancerous tissue sections, to other biological domains (e.g., dermatology), as well as the extent to which prediction modeling can preserve salient biological pathways and relationships required for downstream analysis on larger cohorts.

In this pilot study, we develop and validate a deep learning method for the prediction of spatial gene expression across spatially variable genes in routine H&E-stained skin tissue. Predictions can be used to create synthetic multidimensional tissue maps—similar to those produced through spatial transcriptomic profiling—for tissues without corresponding spatial transcriptomics data. Use of deep learning models promises to reduce the cost and time associated with spatial transcriptomics data acquisition for dermatological applications, greatly expanding access to the technology and its range of unique insights. Interrogating pathways associated with spatially inferred genes can advance our knowledge of skin biology, improve diagnostic tools, and pave the way for more personalized treatment strategies.

## 2. Methods and Materials

In this work, we attempt to predict the spatial gene expression of Visium spatial transcriptomics spots distributed across 40X magnification H&E slides. To this end, we use the following methods:

1. **Data collection and annotation:** Acquired H&E whole slide images (WSI), and spatially registered Visium CytAssist assayed spatial transcriptomics slides from 4 human cheek skin tissue samples collected from sites histologically adjacent to basal cell carcinoma (BCC) and squamous cell carcinoma (SCC) during skin cancer removal surgery. These samples were then graded by dermatologists for their solar elastosis status (two mild, two severe). Additionally, dermatologists annotated regions of each corresponding to distinct histological entities (e.g., epidermis, eccrine glands, hair follicles, sebaceous glands, and vascular/endothelial infrastructure).
2. **Preprocessing:** Preprocess gene expression and WSI subarrays to capture spatially variable genes and genetically dense regions of tissue.
3. **Model development:** Configure the SWIN-T transformer to perform two distinct modeling tasks, binary (dichotomized expression) and continuous gene expression prediction, on the 1000 most spatially variable genes.
4. **Leave one-patient-out cross-validation:** Evaluation on held-out slides/patients as a measure of external applicability.
5. **Recover spatial biology inferences:** Model performance was further measured using: 1) pathway analysis for high performing genes, 2) topological consistency between ground truth and predicted expression and 3) the ability to recapitulate genes and pathways associated with distinct histological structures.

Each of these steps will be further detailed in the ensuing sections.

### 2.1. Data Collection

Four specimens were collected for profiling from a cohort of 261 tissue samples obtained in a single site Mohs micrographic surgery (MMS) clinic between March 1^st^ 2022 and October 10^th^ 2022. The samples were mostly from the head and neck, and all from sites histologically adjacent to either basal cell carcinoma or squamous cell carcinoma, as confirmed by histologic analysis of frozen section slides. The tissue was removed as part of standard surgical practice as Burow’s triangle flaps for skin grafting/reconstruction. Triangles are normally discarded—two triangles were collected per patient, in some cases bisected. One triangle underwent formalin fixation while the other triangle was frozen. Formalin fixed specimens were breadloafed, encased in paraffin embedded tissue blocks, and sectioned and stained for hematoxylin and eosin (H&E) using Autostainers for subsequent imaging at 40x resolution (0.25 micron/pixel) using Aperio GT450 scanners. Tissue slides were transported to the Genomics core, where after tissue decoverslipping, the Visium CytAssist device was used to transfer transcriptomic probes from the original glass slides to 11mmx11mm capture areas on Visium slides. Sections from two patients were placed into each capture area to conserve costs and separated during the analysis stage. Whole transcriptomic profiling was accomplished after mRNA permeabilization, poly(A) capture and probe hybridization.

The eosin stain for the tissue sections were imaged using CytAssist, which were then co-registered to the original 40X whole slide images (WSI). Given the limited sample size due to the spatial transcriptomics assay costs, four specimens were selected, representing cheek tissue from four females, two with mild elastosis (participants 178 and 14, ages 24 and 76 respectively), two with severe elastosis (participants 167 and 107, ages 55 and 84) respectively.

### 2.2. Preprocessing

Prior to processing, Visium spatial transcriptomics profiles for samples contained 18,085 genes measured across several thousand locations throughout each slide. Each profile was then subjected to preliminary filtering, where genes and spots were filtered according to their abundance (i.e., cells with less than 500 genes, genes expressed in less than 3 cells, and cells with more than 15% mitochondrial gene expression were filtered out). After filtering out the regions lacking tissue using a custom annotation tool augmented by the SAM, the total number of Visium spots per slide reached 2561, 3279, 3547, and 1737, each sampled in a honeycomb formation. Each Visium spot covers a circular capture area with a diameter of 50-micron (∼200 pixels) at 40x magnification. After sequencing, we used the SpaceRanger package to preprocess the Visium reads into gene count matrices.

Every whole slide image (WSIs) used for the Visium assay captures an area (size of capture area– 11 × 5.5 mm– half the capture area per patient) that spans tens of thousands of pixels along each dimension. Accordingly, to make the prediction task computationally tractable, we subdivided every WSI into square 512 by 512-pixel image patches (i.e., subarrays) centered on each Visium spot. The gene expression of the central 50-micron Visium spots were aligned to each image patch. Data present within the image patch but falling outside the capture area of the Visium spot were considered to have less direct relevance to the cells being assayed. Spots were additionally annotated based on the aforementioned tissue histological structure using the Annotorious OpenSeadragon plugin.

### 2.3. Model Development

#### 2.3.1. Inference Targets

As predicting all of the genes assayed is computationally intractable, we used the SpatialDE library to select the top 1000 genes based on their mean spatial variance (MSV) across all slides (i.e., selected genes that exhibited the greatest spatial variation across the 4 slides). We then tested the capacity of our models to predict both dichotomized and log gene expression for all 1000 genes.

More specifically, in the dichotomized prediction task, patches were classified as having a “high” or “low” gene expression for each gene if the expression of the gene at that patch location was greater or lower than its mean gene expression across all other Visium spots in the corresponding WSI. This approach follows existing work detailed in Fatemi et al.^5^ For this task, models were trained using a binary cross entropy loss function.

In the continuous expression task, by contrast, models were trained to predict the log-pseudocount log(1+counts) gene expression for each gene within the corresponding image patch region. For this task, loss was calculated using the mean squared error, which was found to comparable to modeling counts using the zero-inflated negative binomial distribution.^5^

#### 2.3.2. Modeling Approach

Previous work has established the importance of spatial and neighborhood context information in both the dichotomized and continuous gene expression tasks.^5^ In this study, we leveraged the SWIN-T vision transformer, a hierarchical transformer that has gained repute for building hierarchical feature maps by iteratively merging information from nearby image patches in deeper layers.^7^ Transformers divide images into smaller subimages, and numerical descriptors are extracted for each subimage using convolutional filters, along with information on the relative positioning of the subimages. Self-attention mechanisms are used to route information across the image based the relevance of one subregion of the image to another. In both the dichotomized and continuous expression tasks, the output layer of the base SWIN-T model was modified. In particular, the output layer was expanded to consist of two feed-forward layers of sizes 768 and 2000, chosen through coarse experimentation to maximize model performance. Both the dichotomized and continuous expression models yielded predictions for the 1000 most spatially variable genes.

#### 2.3.3. Data Augmentation and Hyperparameter Selection

To improve the robustness and generalizability of these models to varied histological contexts, all images in the training set were subject to a series of data augmentation transformations implemented using the Albumentations package.^8^ Images were first resized to 448 by 448 pixels in size, the input dimensionality for the SWIN-T model. Horizontal flips and random brightness contrast were then performed with probabilities 0.5 and 0.2, respectively. A shift, scale, and rotate transformation was also applied to every image with a probability of 0.3. A shifting limit of 0.1, a scaling limit of 0.1, and a rotation limit of 30 were used here. Additionally, random rectangular areas of the images were erased– a maximum of 8 holes were produced per image, each hole obscuring at most 16 by 16 pixels.

Hyperparameters were obtained for both the dichotomized and continuous expression models via a coarse hyperparameter grid search. For the dichotomized models, optimal performance was observed while using a batch size, learning rate, and training length of 64, 0.5×10−6, and 20 epochs. Whereas for the continuous models, optimal performance was observed while using a batch size, learning rate, and training length of 64, 0.33 × 10−6, and 20 epochs. The Lion optimizer was used in both cases.^9^

### 2.4. Cross Validation

Model performance was measured via leave-one-patient-out cross-validation (LOOCV). In this procedure, three of the four Visium spatial transcriptomics samples were used for training and validation, while the remaining sample was used for testing. This procedure was repeated four times to account for all possible training/testing combinations. Reported performance metrics for each gene are the macro-averaged (across slides, weighting each slide equally) median (across genes) area under the receiver operating characteristic curve (AUROC) and F1 score (F1) statistics for the dichotomized task, and correlation coefficients (Spearman coefficient) to compare true versus predicted pseudocounts– log(1+counts)– for the continuous task. Macro performance statistics underwent 1,000 sample nonparametric bootstrapping across the Visium spots to yield 95% confidence intervals.

### 2.5. Biological Salience

To assess for the model’s ability to capture meaningful biological information from tissue histology, model predictions were also scrutinized for their ability to 1) recapitulate a range of biologically salient pathways, 2) maintain the shape and spatial signature of ground truth spatial gene expression data in lower dimensional space (i.e., preserves key relationships; spots cluster similarly) using the aligned-UMAP procedure, and 3) facilitate the inference of biologically salient features, such as histological markers. These tasks are detailed below.

#### 2.5.1. Pathway Analyses

Given the nature of histomorphological data, it is unreasonable to expect that every gene can be predicted from tissue histology alone. Accordingly, we sought to determine the biological pathways associated with sets of differentially performing genes to answer understand what biological properties make a gene amenable to prediction. We utilized the GO Biological Process 2023 database through the EnrichR package, to perform a pathway analysis on predicted genes stratified by decile after ranking genes based on predictive performance.^10, 11^ The top 3 pathways were selected for each decile—from 90^th^ to 100^th^ decile (i.e., top performing genes) to the 0^th^ to 10^th^ device (i.e., worst performing genes)—based on their combined score (i.e., magnitude of representation and statistical significance). Detected pathways were also filtered by tissue specificity (i.e., could reasonably be involved with the skin).

We further sought to identify whether the gene signatures correspondent to different histological architectures was congruent between true and predicted expression. First, the top 100 most differentially expressed genes were found using the Wilcoxon rank-sum test in a one vs. rest fashion for each tissue architecture (e.g., follicles versus non-follicular structures) using both predicted and ground truth data. A pathway analysis using GO Biological Process 2023 database through the EnrichR package was then performed for the top true and predicted differentially expressed genes for each architecture. The top 10 pathways were selected by combined score for each histological category. Detected pathways were compared between ground truth and predicted gene expression for each sample under the hypothesis that similar pathways should be associated with the same architectures.

#### 2.5.2. Similar Clustering of Visium Spots and Consistent Topology via Aligned-UMAP

Model predictions were further assessed for their ability recapitulate the topology (i.e., relationships between spots) of ground truth Visium spatial gene expression data within a lower dimensional space. This was accomplished through the comparison of Uniform Manifold Approximation and Projection (UMAP) embeddings (i.e., numerical representations that could be plotted in a 2D scatterplot; closer points share similar expression/biological relevance) for the ground truth and predicted expression profiles (on held-out slides) extracted using the SWIN-T model. Ground truth and predicted gene expression profiles for each slide were co-projected to a lower dimensional space using the Aligned-UMAP procedure to preserve the relative orientation and alignment between spots to enable comparison between the approaches. Each Visium spot from the WSI was plotted as 2D scatterplot point and colored according to its gene expression profile as dictated by the Leiden clustering algorithm. In other words, ground truth Visium spots sharing similar transcriptional information are grouped to the same Leiden cluster, while genetically dissimilar spots are grouped to different Leiden clusters. These ground truth cluster assignments were overlaid on the scatterplots for the predicted expression patterns. It is expected that the relative positioning between the clusters would be preserved in the 2D scatterplot for the predicted expression, which would measure the extent to which model predictions recapitulated patterns associated with distinct histological regions of each WSI.

Aside from overlaying the original ground truth clusters, predicted expression profiles were also separately clustered through the Leiden algorithm, yielding a separate set of cluster assignments for the same Visium spots. These assignments were compared to the ground truth Visium spatial transcriptomics profiles’ clusters. Similar clustering assignments would provide further evidence for greater correspondence between transcriptional data and information derived from the histology.

## 3. Results^†^

### 3.1. Prediction of Spatial Transcriptomic Patterns from Histology

In the dichotomized prediction task, the SWIN-T vision transformer model achieved a macro-averaged (i.e., across genes) median AUC and F1 score of 0.80 and 0.61, respectively, across the testing sets (**Supplementary Table 1**). The model performed best on genes *ADIPOQ*, *PLIN1*, and *PKP3* (involved in fatty acid metabolism^12^, triacylglycerol storage^13^, and desmosome function and stability^14^, respectively), and worst on genes *ANKRD35*, *ALAS1*, and *MIA* (of which the latter two are known to be involved in heme biosynthesis^15^ and melanocyte migration^16^, respectively). Dichotomized model predictions for genes *ADIPOQ*, *PLIN1*, and *PKP3* are visualized across samples 14 and 178 (**Figure 2**), demonstrating spatial concordance between true and predicted expression. In the continuous prediction task, models achieved a macro-averaged median Spearman coefficient of 0.60 across the testing sets (**Supplementary Table 1**). The model performed best on genes *KRT14*, *CXCL14*, and *COL1A2* (involved in epithelial cell integrity^17^, keratinocyte function^18^, and collagen synthesis^19^, respectively) and worst on genes *CKM*, *MYLPF*, and *ODF21* (the former two are known to be involved in energy homeostasis^20^ and muscle development^21^). Continuous model predictions for genes *KRT14*, *CXCL14*, *PI16* were visualized across samples 107 and 167 (**Supplementary Figure 2**), demonstrating spatial concordance between true and predicted expression. Note that models in both the dichotomized and continuous prediction tasks were trained to predict the same set of 1000 spatially variable genes.

**Figure 1:**
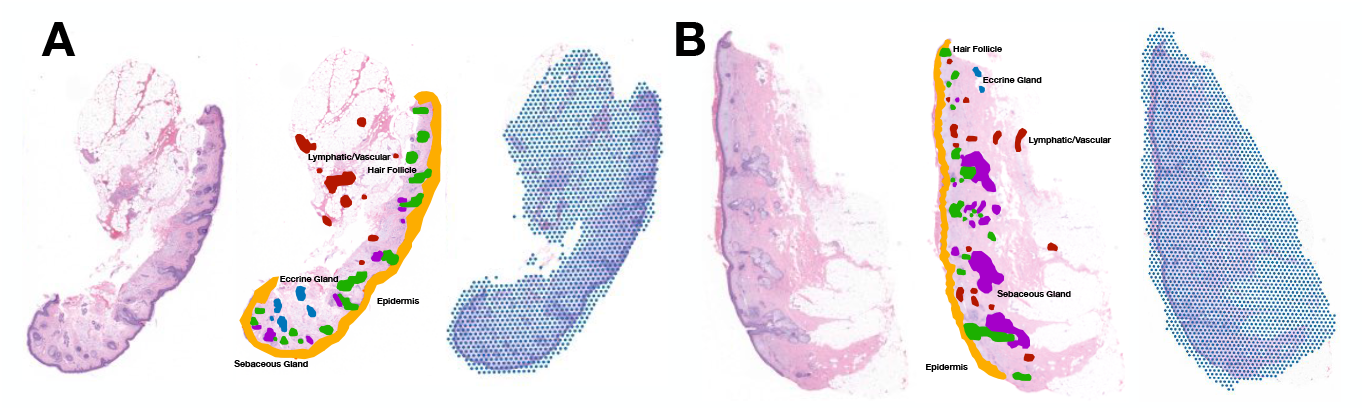
Cohort Description. 261 WSIs were scanned. Four of these slides underwent further spatial transcriptomics profiling and were annotated for distinct histological architectures. Two are shown here. From left to right, the WSI, histological annotations, and Visium spatial transcriptomics spot array for **(A)** sample 14 and **(B)** sample 167.

**Figure 2:**
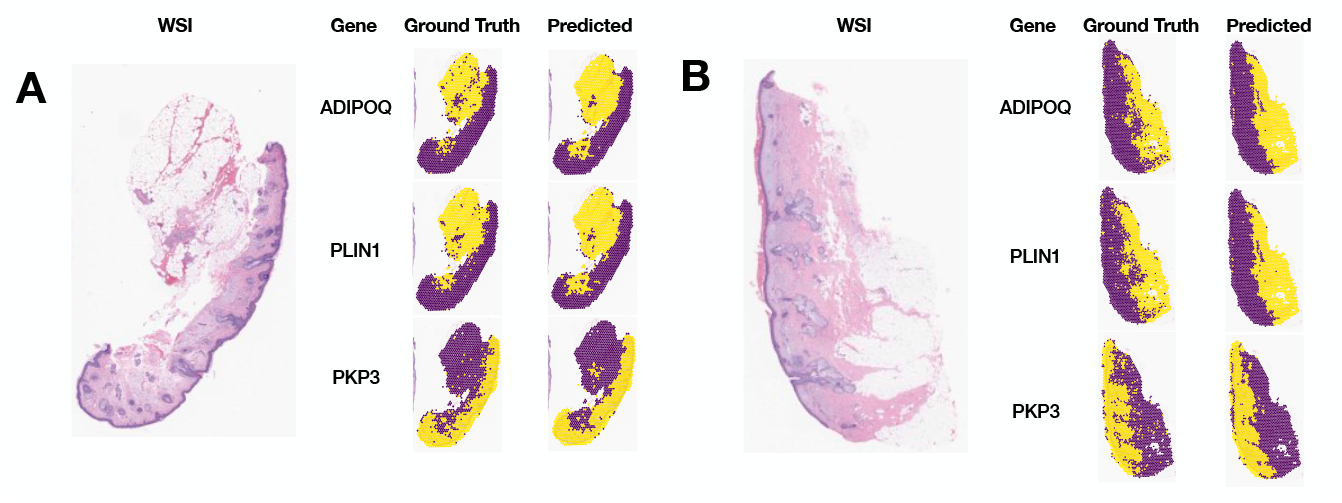
Dichotomized RNA Expression Prediction. Dichotomized spatial gene expression was inferred for **(A)** samples 14 and **(B)** 167 and compared with the respective ground truths. A spot is colored yellow if gene expression in this spot exceeds global mean gene expression. Performance is displayed for the top performing genes, ADIPOQ, PLIN1, and PKP3, which achieved macro-averaged AUC values of 0.942, 0.938, and 0.918, respectively.

### 3.2. Pathway Analysis

For each performance decile, the top 3 most salient biological pathways by combined score were determined (**Supplementary Table 2**). Across both the dichotomized and continuous prediction tasks, biological pathways associated with the top performance decile (i.e., 90^th^ to 100^th^ percentile genes ranked by performance) pertained to skin and epidermis development and maintenance, skin cell proliferation, and the regulation of extracellular matrix and cell-cell adhesion (**Table 1**). By contrast, biological pathways associated with genes in the worst performance decile (i.e., 0^th^ to 10^th^ percentile genes ranked by performance) across both the dichotomized and continuous prediction were far less associated with relevant biological phenomena, pertaining to immune signaling, cell-turnover regulation, gas transport, and muscle cell development (**Table 1**). More generally, biological pathways associated with higher performing genes tended to be more closely related to skin development, differentiation, pigmentation, and fat metabolism, while distinct trends were less clear for those biological pathways associated with lower performing genes (**Supplementary Table 2**).

**Table 1:** Performance Pathway Analysis. Combined performance statistics for both the dichotomized and continuous models were used to perform a performance-stratified pathway analysis. AUC and Spearman coefficient were used to stratify genes in the dichotomized and continuous tasks, respectively. The top 3 pathways, measured via EnrichR using the Go Biological Process 2023 database, are reported for the highest and lowest performance deciles. Refer to **Supplementary Table 2** for an extended version of this table detailing all performance deciles.

| Gene Performance | Task | Pathway | Score | Overlap | P-value |
| --- | --- | --- | --- | --- | --- |
| **Top performing genes: 90-100<sup>th</sup> Percentile Genes** | **Dichotomized** | Establishment of Skin Barrier | 2127 | 6/19 | 6.9E-10 |
|  |  | Skin Epidermis Development | 1785 | 6/21 | 4.0E-04 |
|  |  | Keratinocyte Proliferation | 803 | 2/6 | 1.4E-11 |
|  | **Continuous** | Positive Regulation of Epidermis Development | 2725 | 4/9 | 7.3E-08 |
|  |  | Desmosome Organization | 2654 | 3/6 | 2.4E-06 |
|  |  | Intermediate Filament Bundle Assembly | 1905 | 4/7 | 4.2E-06 |
| **Bottom performing genes: 0-10<sup>th</sup> Percentile Genes** | **Dichotomized** | Interleukin-2-Mediated Signaling Pathway | 615 | 2/7 | 2.8E-04 |
|  |  | Cellular Response to Interleukin-2 | 615 | 2/7 | 5.1E-04 |
|  |  | Negative Regulation of T Cell Apoptotic Process | 615 | 2/7 | 5.1E-04 |
|  | **Continuous** | Gas Transport | 504 | 3/15 | 5.3E-05 |
|  |  | Positive Regulation of Respiratory Burst | 493 | 2/8 | 6.7E-04 |
|  |  | Regulation Of Skeletal Muscle Cell Differentiation | 493 | 2/8 | 6.7E-04 |

The top 100 differentially expressed genes for each histological sub-type were determined for both ground truth and predicted data samples, and these genes were leveraged for further pathway analyses. The pathway analysis results for the top 100 differentially expressed genes for dichotomized and continuous gene expression data are reported in **Supplementary Table 3** and **Supplementary Table 4**. Across both dichotomized and continuous gene expression data, pathways associated with the formation of the sebaceous gland and epidermis were found to be in high agreement between ground truth and predicted expression, while the agreement was more modest other in histological sub-types (**Table 1; Supplementary Table 3**).

### 3.3. Topological Consistency

#### 3.3.1. Leiden Clustering

A visual inspection of the aligned-UMAP diagrams demonstrates similar clustering patterns and topological consistency between the predicted and the ground truth expression data across both models trained for dichotomized and continuous regression tasks (**Figure 3; Supplementary Figure 2**). We noted that the Leiden clusters assigned to the ground truth expression were similar to those assigned to predicted expression embeddings. Nonetheless, differences remained. We did not observe complete separation in the predicted expression embeddings, representing a fuzzier or more connected/intermediate topological structure (**Figure 3; Supplementary Figure 2)**. These spots in the predicted data were, accordingly, located between Leiden clusters more often than spots in the ground truth genetic data, where Leiden clusters tended to be far more spatially distinct. This feature was noted for both dichotomized and continuous expression models, although this pattern was more prevalent for dichotomized expression (**Figure 3; Supplementary Figure 2)**.

**Figure 3:**
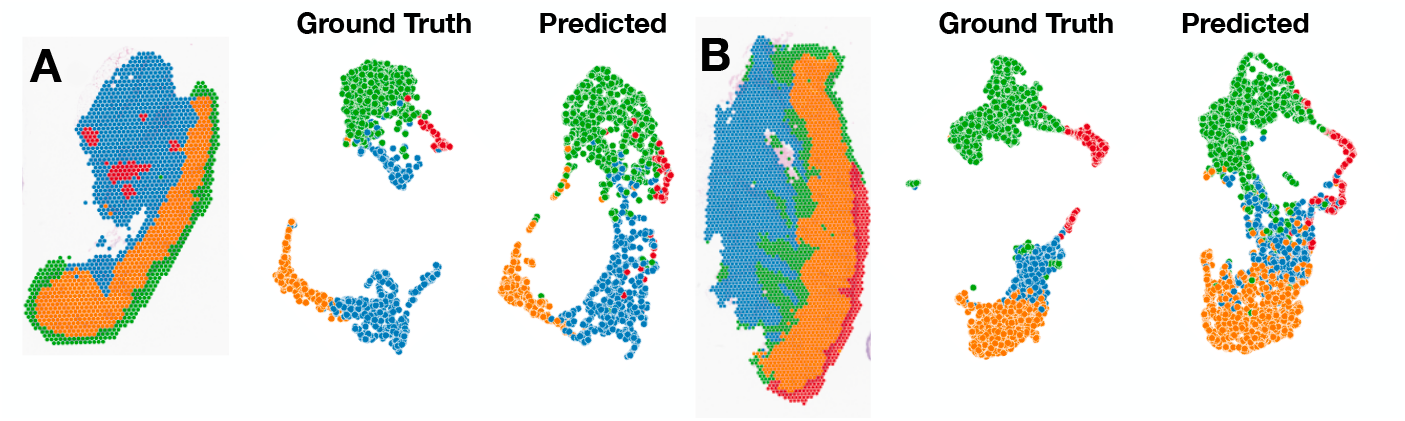
Dichotomized Expression Topological Analysis. **(A)** From left to right, ground truth spatial Leiden clustering, ground truth aligned UMAP, and predicted aligned UMAP for sample 14. **(B)** From left to right, ground truth spatial Leiden clustering, ground truth aligned UMAP, and predicted aligned UMAP for sample 178. In both rows, the same spots are colored identically according to their ground truth gene expression profiles after Leiden clustering analysis. Dichotomized gene expression data was used here. The Leiden clustering resolution was set to 0.2.

Model predictions in both the dichotomized and continuous expression tasks also preserved the general shape of ground truth genetic data while plotted across each whole slide image (**Supplementary Figures 3** and **4**). We further observed, however, that predicted data tended to contain genetically intermediate states, as evidenced by the greater number of Leiden clusters produced in the predicted data compared to the ground truth data while using the same Leiden clustering resolution (**Supplementary Figures 3** and **4**). Though models in both tasks produced data that captured larger macro-architectural differences in gene expression found across skin tissue, the dichotomized model tended to produce data that more closely preserved the relationships determined using Leiden clustering plotted across the slide found across the ground truth data (**Supplementary Figure 3**). Models in the continuous expression task, though high performing, tended to produce data that recapitulated the spatial genetic variation of macro-architectural features in skin tissue less well, evidenced, again, by disparities in the number and placement of Leiden clusters when comparing the predicted and ground truth data (**Supplementary Figure 4**).

#### 3.3.2. Histological Annotations

Performing Aligned-UMAP on the ground truth and predicted expression data for Visium spots tagged by histological structures demonstrated that embeddings in both groups clustered by distinct histological regions of skin tissue (**Figure 4; Supplementary Figure 5**). That is, Visium spots corresponding to similar histological structures clustered in similar locations across both UMAP plots, preserving the genetic relationships between these histological architectures. The distinctness of these clusters was preserved for both dichotomized and continuous gene expression predictions, though predicted continuous expression data appeared to preserve the topology better than dichotomized gene expression data (**Figure 4; Supplementary Figure 5**).

**Figure 4:**
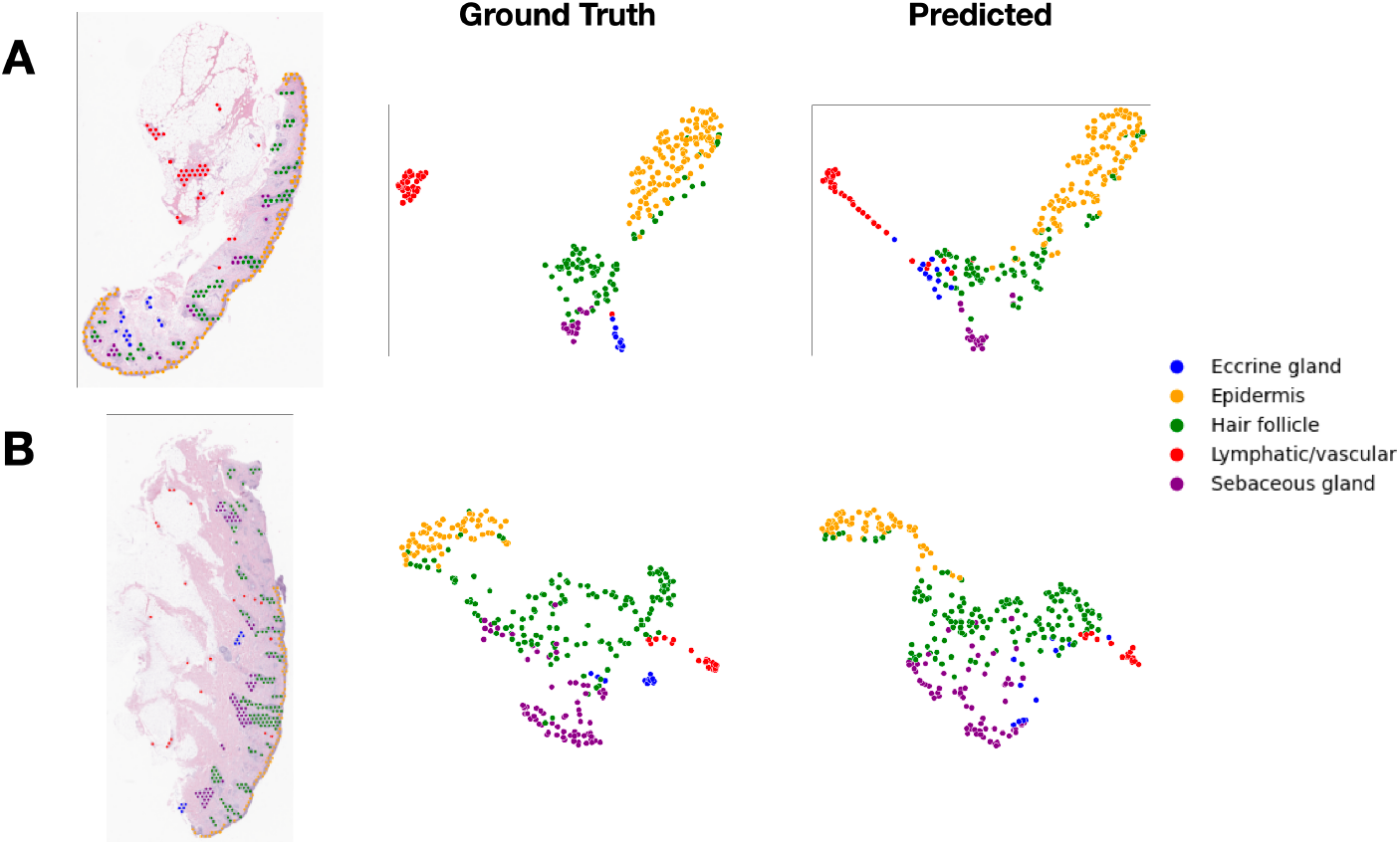
Dichotomized Expression Histological Analysis. Aligned-UMAP procedure was used to reduce the dimensionality of both the ground truth and predicted dichotomized gene expression vectors for **(A)** sample 14 and **(B)** sample 178. Spots are colored according to their histological annotations.

## 4. Discussion

In this work, we developed a set of spatial gene expression inference models for histopathologically normal skin tissue in the context of molecular changes associated with photoaging. We make use of the novel CytAssist co-registration/imaging technology, allowing for 40X resolution imaging of tissue slides. Beyond quantitative validation of performance (e.g., AUC, F1, Spearman coefficient), we also reaffirmed the biological relevance of the predicted expression pattern. In particular, extracted histological features from our models remained faithful to underlying biological pathways, buttressing their potential use across a range of biological inference tasks, and lending credibility to their role in democratizing the spatial transcriptomics paradigm to the broader research community.

With the Visium CytAssist technology, our models were trained with histological information at >4 times the spatial resolution of previous studies. We achieved comparable performance to a prior study that utilized an Inception convolutional neural network for both dichotomized and continuous prediction of gene expression.^5^ While acknowledging the need for caution when comparing these models, as they represent different biological domains (Skin vs Colon; different genes used in the models), observed performance disparities may arise from variations in methodology. These differences could include the utilization of distinct imaging resolutions (40X vs 20X) or the selection of different modeling approaches (ZINB vs raw log gene expression). Since our predictions were made within a more focused visual receptive field, disregarding the surrounding wider tissue architecture, future work can explore the examination of larger-scale histological context.

The pathway and topological analysis provided useful insights on the nature of spatial RNA inference from histology. It is important to highlight that our findings suggest that genes with a clear histological basis are more likely to be accurately predicted compared to genes lacking such a theoretical histological foundation. Our results also demonstrate the importance of developing models germane to the biological question at hand: a model trained on colon tissue is not expected to perform well on skin tissue. Hence, investigating modeling approaches that prioritize specific biological phenomena emerges as a promising direction for future research. Genes that demonstrate good performance, or effectively recapitulate histological patterns, could potentially be utilized for further research applications across larger cohorts.

The topological analysis demonstrated that predicted expression profiles did not cluster as distinctly as the original expression patterns. When coloring by Leiden cluster affiliation and histological association, the predicted spot level gene expression fell between ground truth clusters, representing intermediate histological states learned by the neural networks. Future work may seek to understand which histomorphological features relate to different molecular pathways through newly established interpretation approaches^22^. The integration of coregistered slide imaging with spatial molecular information can facilitate such analyses. Moreover, by subsetting predicted and true gene expression based on shared molecular pathways (such as genes involved in epithelium development, cell-cell junctions, immune function, etc.) and conducting comparable topological analyses, it is possible to identify the molecular pathways that exhibit the highest degree of topological distinctiveness. Nevertheless, topological analyses have emerged as a timely and relevant topic in the realm of single-cell and spatial analyses, offering the potential to uncover additional dimensions of cellular and histological heterogeneity^23–25^.

This study reinforces the potential of spatial transcriptomics approaches for research and clinical applications. For example, photoaging, which is linked to skin cancer risk, lacks reliable measurement tools due to variations in histological assessments and self-reported UV exposure. Existing analyses typically focus on specific cellular components (e.g., dermal fibroblasts, elastosis, keratoses), often disregarding or unaware of photoaging-related factors. Expanding spatial molecular findings through RNA inference to a larger cohort can help identify cell-type specific sources of photoaging in specific tissue architectures while controlling for numerous potential confounders and presents an intriguing area of follow up given the models established in this study. Spatial RNA inference can uncover novel cellular components related to precancerous alterations resulting from chronic sun exposure. By targeting profiling of these tissue regions, researchers can explore residual heterogeneity, while examining cell-type specific alterations and additional factors related to accelerated aging, with the caveat that tissue from this cohort is histologically adjacent to surgical site of repair, potentially harboring a cancerization field effect.

In clinical practice, the use of virtual RNA models has the potential to inform treatment planning and assess treatment response. If these models can identify proxy measures of photoaging, spatial molecular inference can be employed to evaluate the effectiveness of skin therapeutics through quantitative assessment of biomolecular changes at screening, baseline, and endpoint. This approach offers a more objective and quantitative measurement of the impact of treatments on skin health and can enhance the validity of therapeutic interventions. Similarly, applications are envisioned for treatment of non-healing skin ulcers and separately hair loss driven by an autoimmune response (e.g., alopecia areata), revealing potential components of relevant immune polarity (e.g., M1/M2 macrophage balance, etc.).^26, 27^ Additionally, virtual RNA inference models can inform disease management options for various solid tumors, functioning similar to immunohistochemical assays (e.g., immunoscore) that shed light on the infiltration of cytotoxic immune cell lineages, identifying independent risk factors of tumor recurrence and survival. Spatial molecular assessments can identify targetable therapeutic pathways for personalized treatment options.

This study is not without limitations that can direct for future research. Our sample size was small, limiting our ability to account for potential variability in histology and surgical sites. Additionally, the non-biopsied nature of the samples and their proximity to potentially precancerous tumor tissue may introduce differences in gene expression related to factors other than UV exposure. Introducing matching normal control tissue, considering factors like limited sun exposure and low field effect potential, along with expanding the cohort to control for additional age ranges, sex and tissue site, could help reveal photoaging differences specific to these groups. Skin tone is another confounding factor that should be addressed, and it can be controlled using measures such as the Fitzpatrick skin phototype scale or derived continuous measures. To improve rigorous quantitative photoaging assessments, various measures of photoaging can be combined using factor analyses, leading to meaningful composite measures, such as DNA methylation age-related measures, elastosis, and UV questionnaires. Addressing these limitations and incorporating a more diverse and extensive sample size can enhance the reliability and applicability of future studies in this field.

## 5. Conclusion

Machine learning technologies that can infer spatial molecular information from routine tissue stains have the potential to facilitate low-cost accessible spatial transcriptomic assessments for large scale molecular epidemiological studies. Such studies can uncover novel risk factors of early photocarcinogenesis or inform relevant treatment/therapeutic options by expanding the set of targetable molecular pathways within specific tissue architectures. Our skin study sets the stage for larger-scale studies to identify spatial molecular correlates of skin sun damage and evaluate novel therapeutics that may reverse this damage. While our models exhibited impressive performance in predicting dichotomized and continuous gene expression within tissue slides, it is crucial to acknowledge the need for further development and validation of this approach. When utilizing these algorithms, it is important to consider the genes that are known to be influenced by histological characteristics. Additionally, any novel findings obtained through these tools should be corroborated and validated using well-established immunostaining techniques, ensuring the reliability and robustness of the results.

## Acknowledgements and Location of Supplementary Materials

The authors acknowledge the support of the Center for Clinical Genomics and Advanced Technology in the Department of Pathology and Laboratory Medicine of the Dartmouth Hitchcock Health System which includes the Pathology Shared Resource, at the Dartmouth Cancer Center with NCI Cancer Center Support Grant 5P30 CA023108-37. Spatial transcriptomics assays were carried out in the Genomics and Molecular Biology Shared Resource (GMBSR) at Dartmouth which is supported by NCI Cancer Center Support Grant 5P30CA023108 and NIH S10 (1S10OD030242) awards. Spatial studies were conducted through the Dartmouth Center for Quantitative Biology in collaboration with the GMBSR with support from NIGMS (P20GM130454) and NIH S10 (S10OD025235) awards. Supplementary materials can be found at the following DOI: https://doi.org/10.5281/zenodo.8197850.

## Supplementary Materials

**Supplementary Figure 1:**
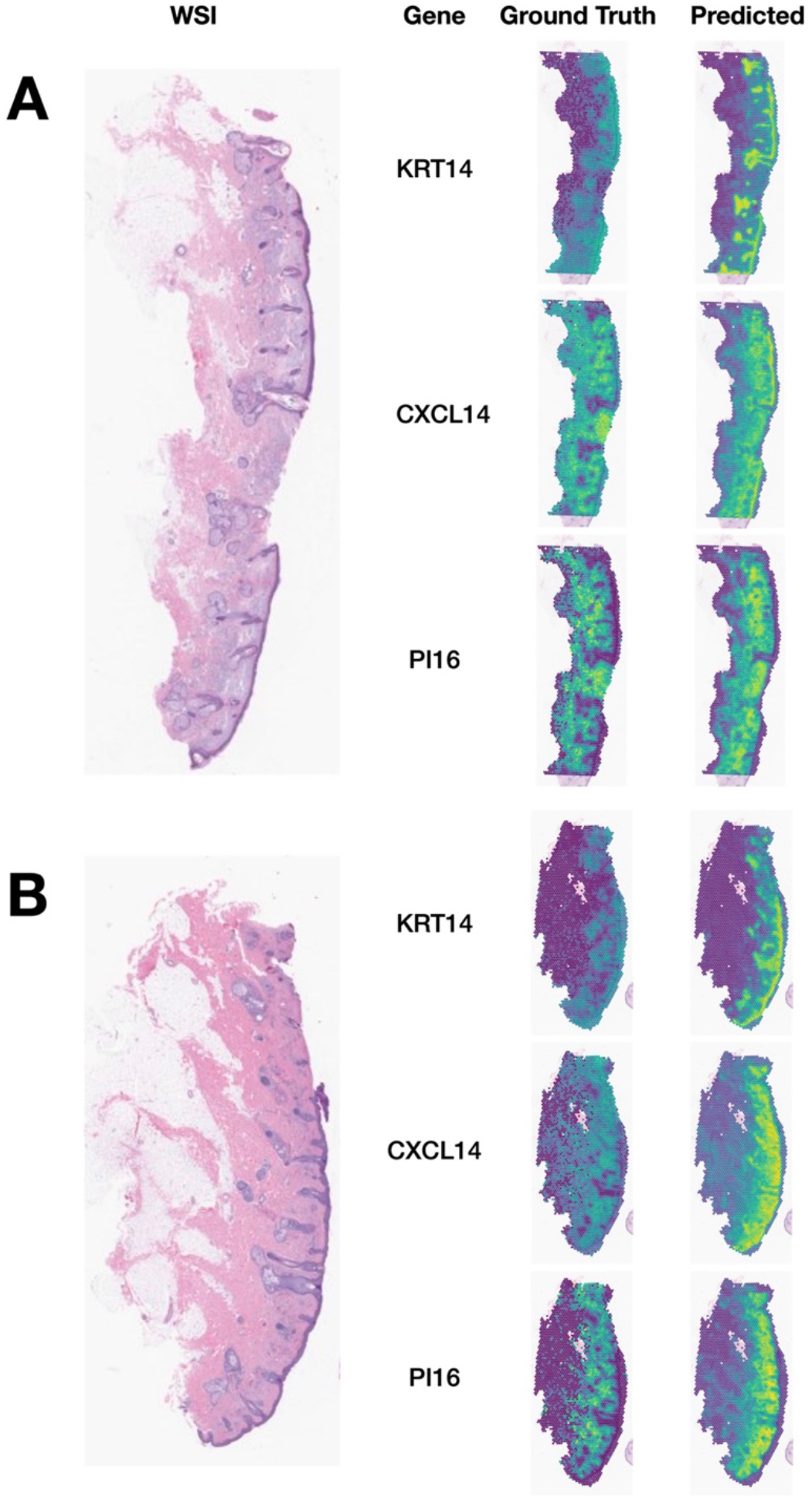
Continuous RNA Expression Prediction. Continuous spatial gene expression was inferred for **(A)** samples 107 and **(B)** 178 and compared with their respective ground truths. The higher gene expression values are displayed in yellow, while lower values are shown in purple. Performance is displayed for several top performing genes, including KRT14, CXCL14, and PI16, which achieved macro-averaged Spearman coefficients of 0.849, 0.846, and 0.822.

**Supplementary Figure 2:**
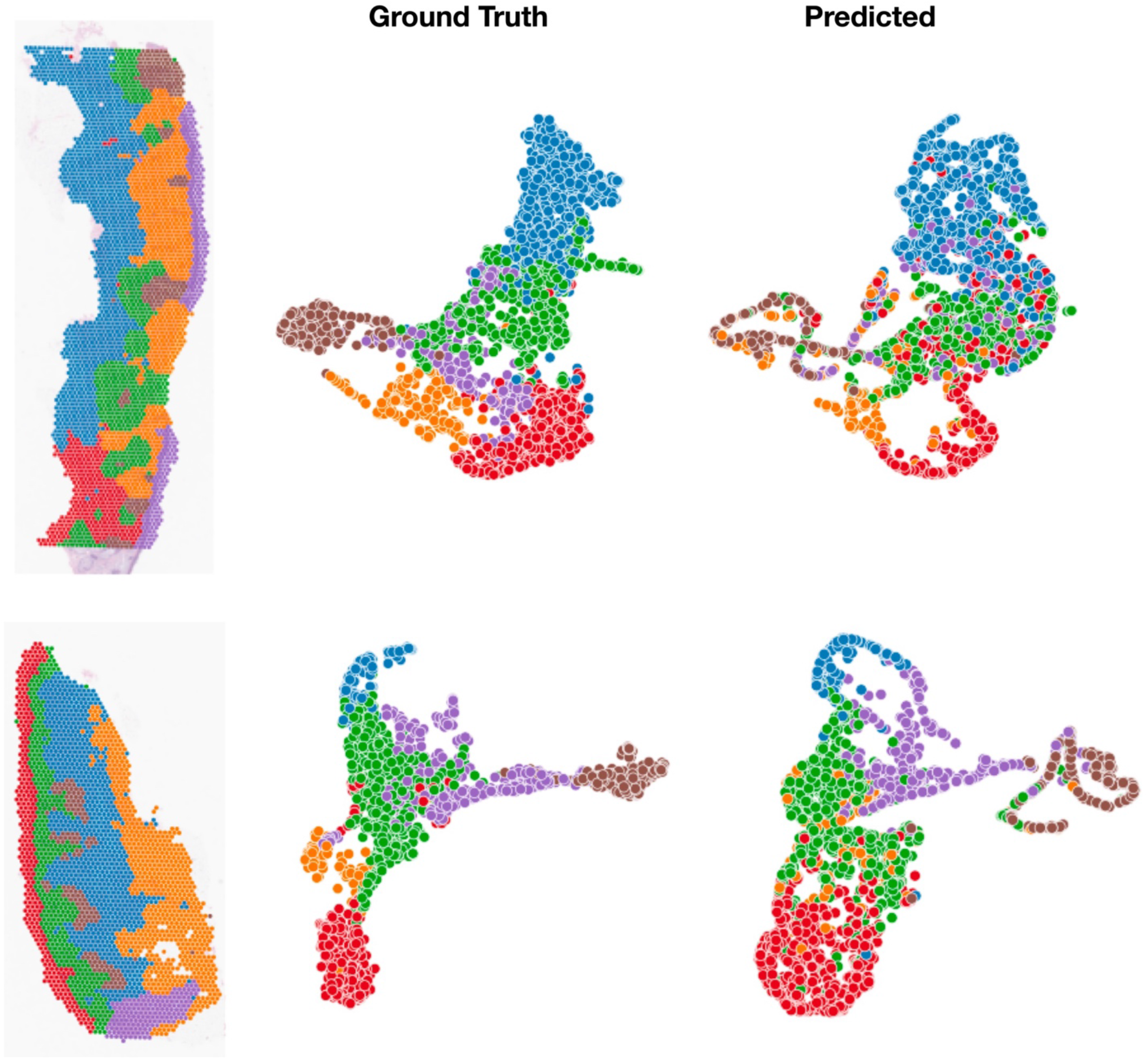
Log Expression Topological Analysis. **(A)** From left to right, ground truth spatial Leiden clustering, ground truth aligned UMAP, and predicted aligned UMAP for sample 107. **(B)** From left to right, ground truth spatial Leiden clustering, ground truth aligned UMAP, and predicted aligned UMAP for sample 167. In both rows, the same spots are colored identically according to their ground truth gene expression profiles after Leiden clustering analysis. Log-transformed gene expression data was used here. The Leiden clustering resolution was set to 0.2.

**Supplementary Figure 3:**
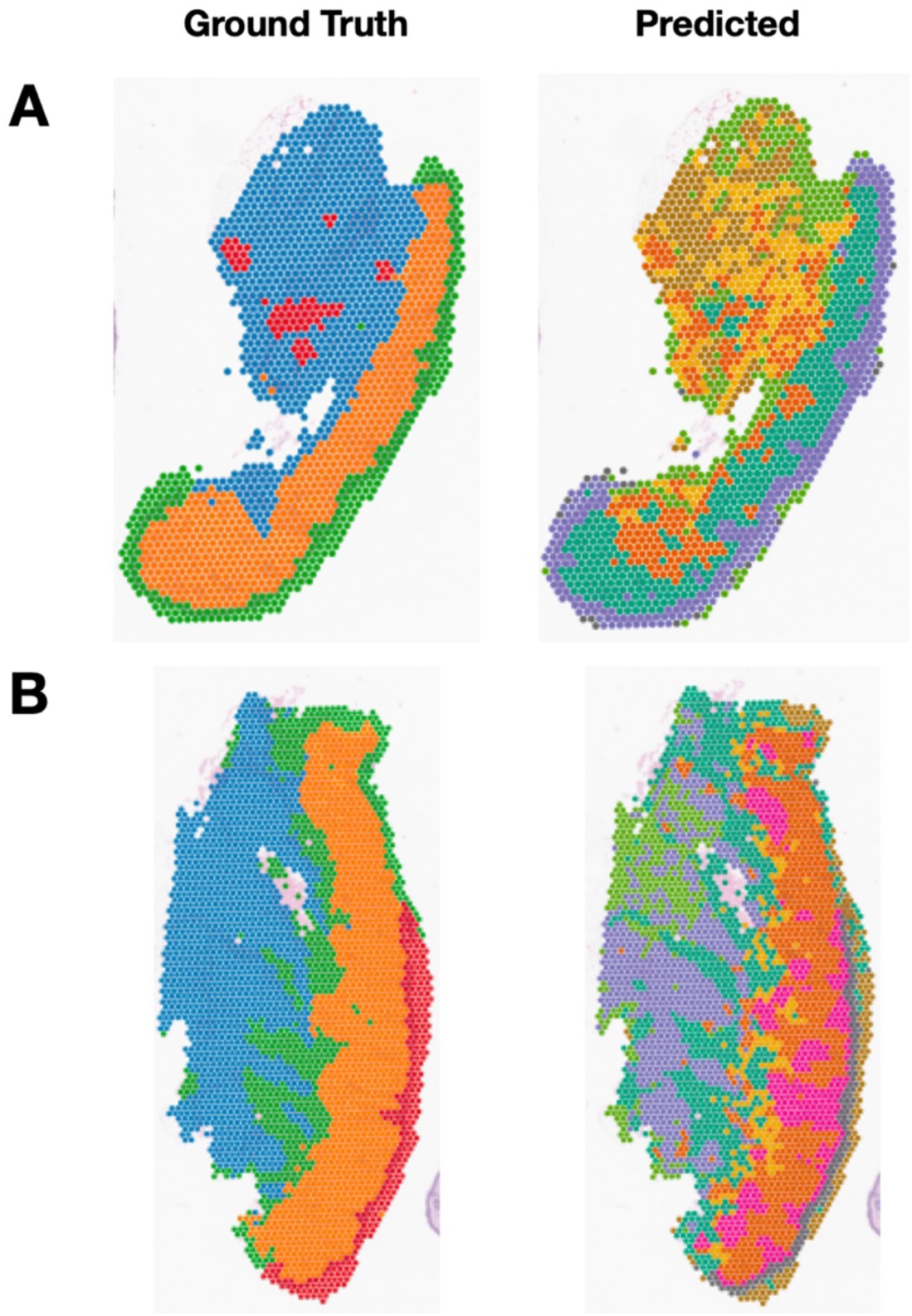
Dichotomized Expression Spatial Leiden Clustering. Visium spots were clustered according to their dichotomized gene expression profiles using both the ground truth and synthetic data for samples **(A)** 14 and **(B)** 178 and visualized spatially. Note that there is no relationship between clusters in the predicted and ground truth cases. A Leiden clustering resolution of 0.2 was used.

**Supplementary Figure 4:**
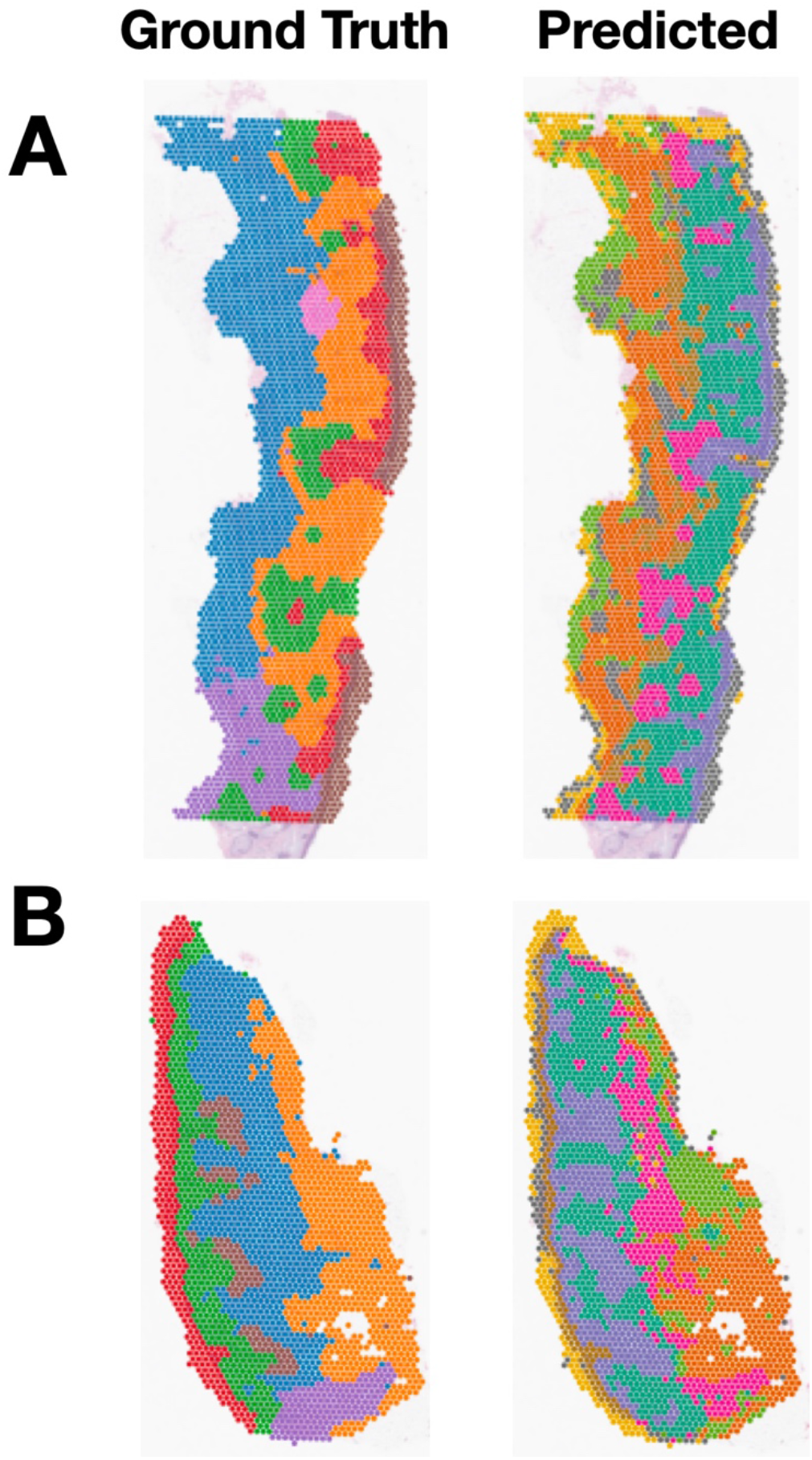
Log Expression Spatial Leiden Clustering. Visium spots were clustered according to their log transformed gene expression profiles using both the ground truth and synthetic data for samples **(A)** 107 and **(B)** 167 and visualized spatially. Note that there is no relationship between clusters in the predicted and ground truth cases. A Leiden clustering resolution of 0.2 was used.

**Supplementary Figure 5:**
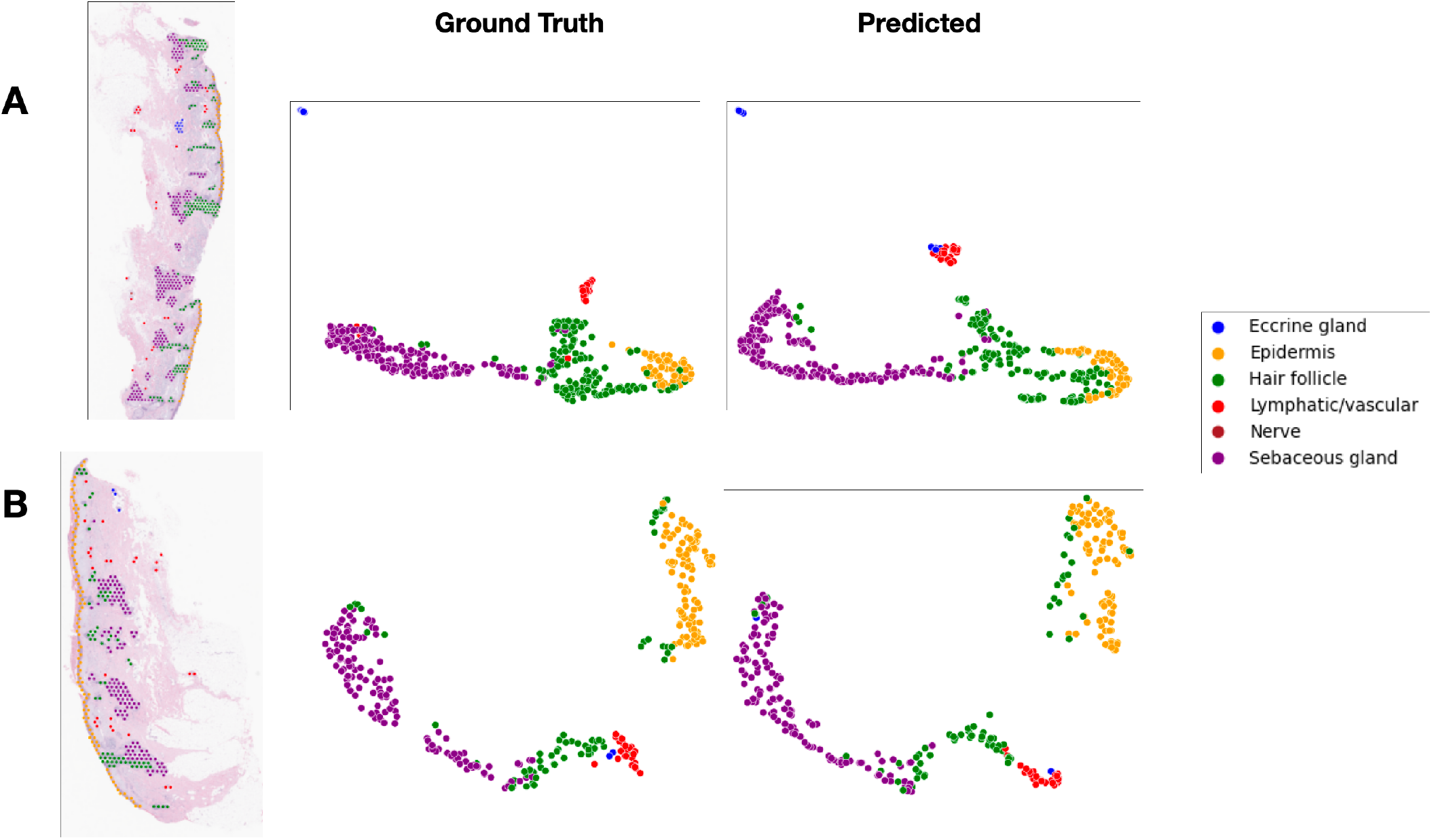
Log Expression Histological Analysis. Aligned-UMAP procedure was used to reduce the dimensionality of both the ground truth and predicted dichotomized gene expression vectors for **(A)** sample 107 and **(B)** sample 167. Spots are colored according to their histological annotations.

**Supplementary Table 1.**
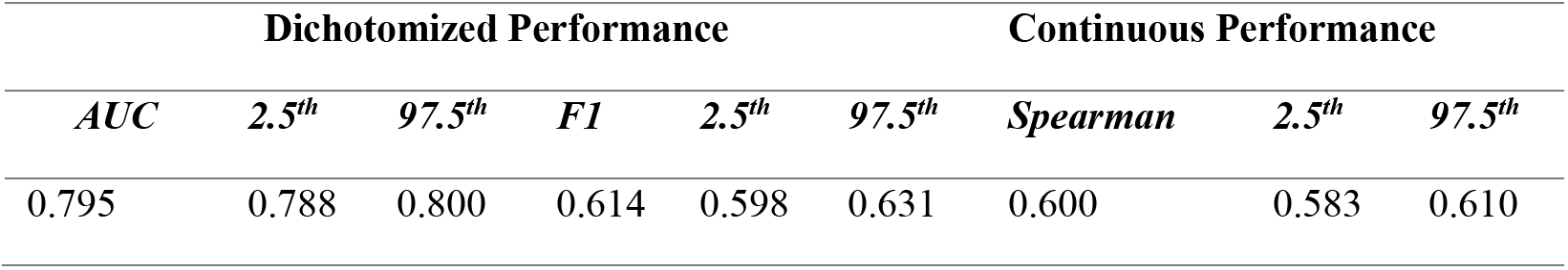
Spatial Gene Expression Prediction Performance. AUC, F1, and Spearman values are the macro-averaged (across slides) median (across genes) performance values. AUC and F1-scores are used to measure the performance of the dichotomized expression models, while the Spearman coefficient was used to measure the performance of the continuous expression model. 95% confidence intervals are reported using 1000-sample non-parametric bootstrapping.

**Supplementary Table 2:**
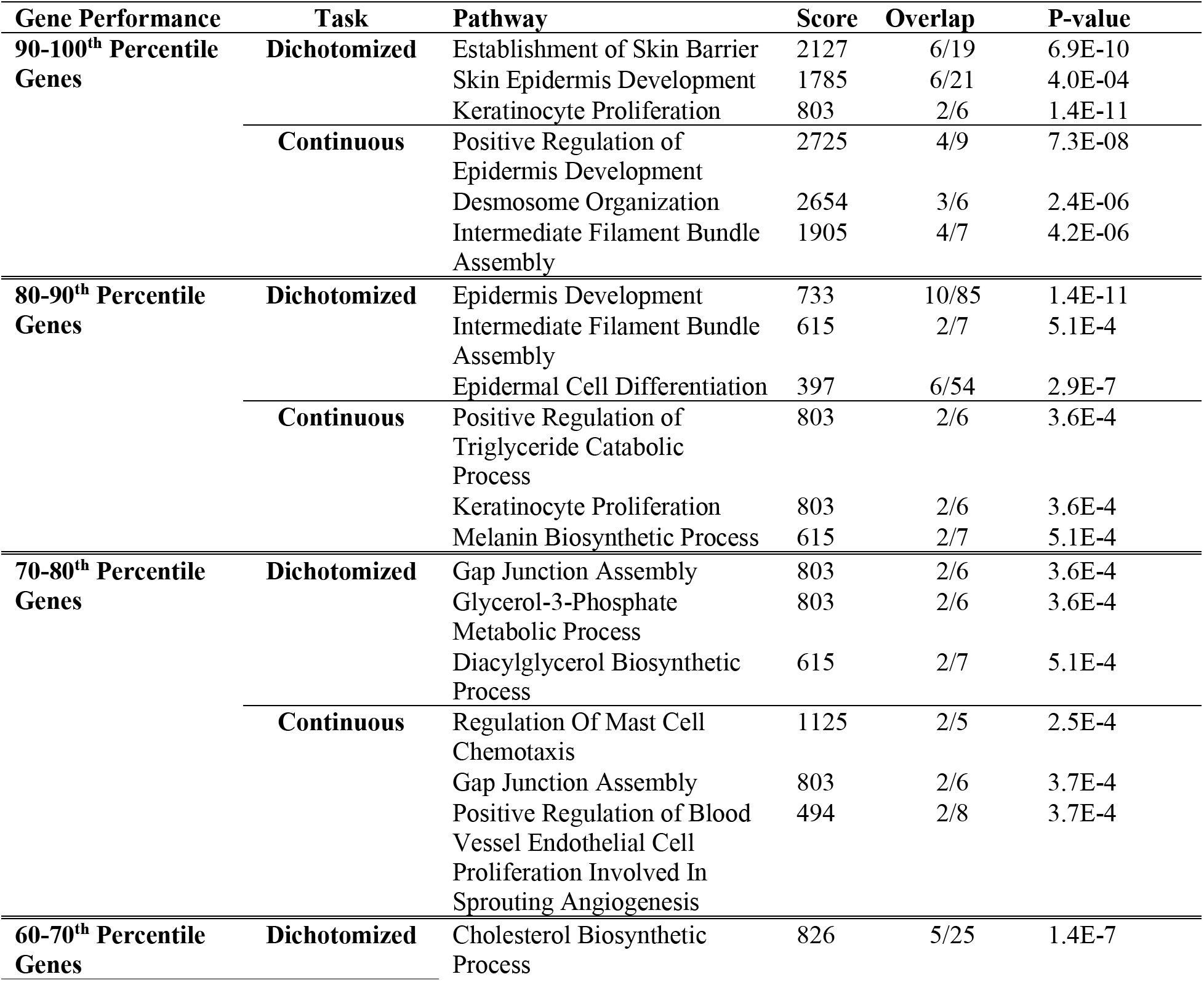
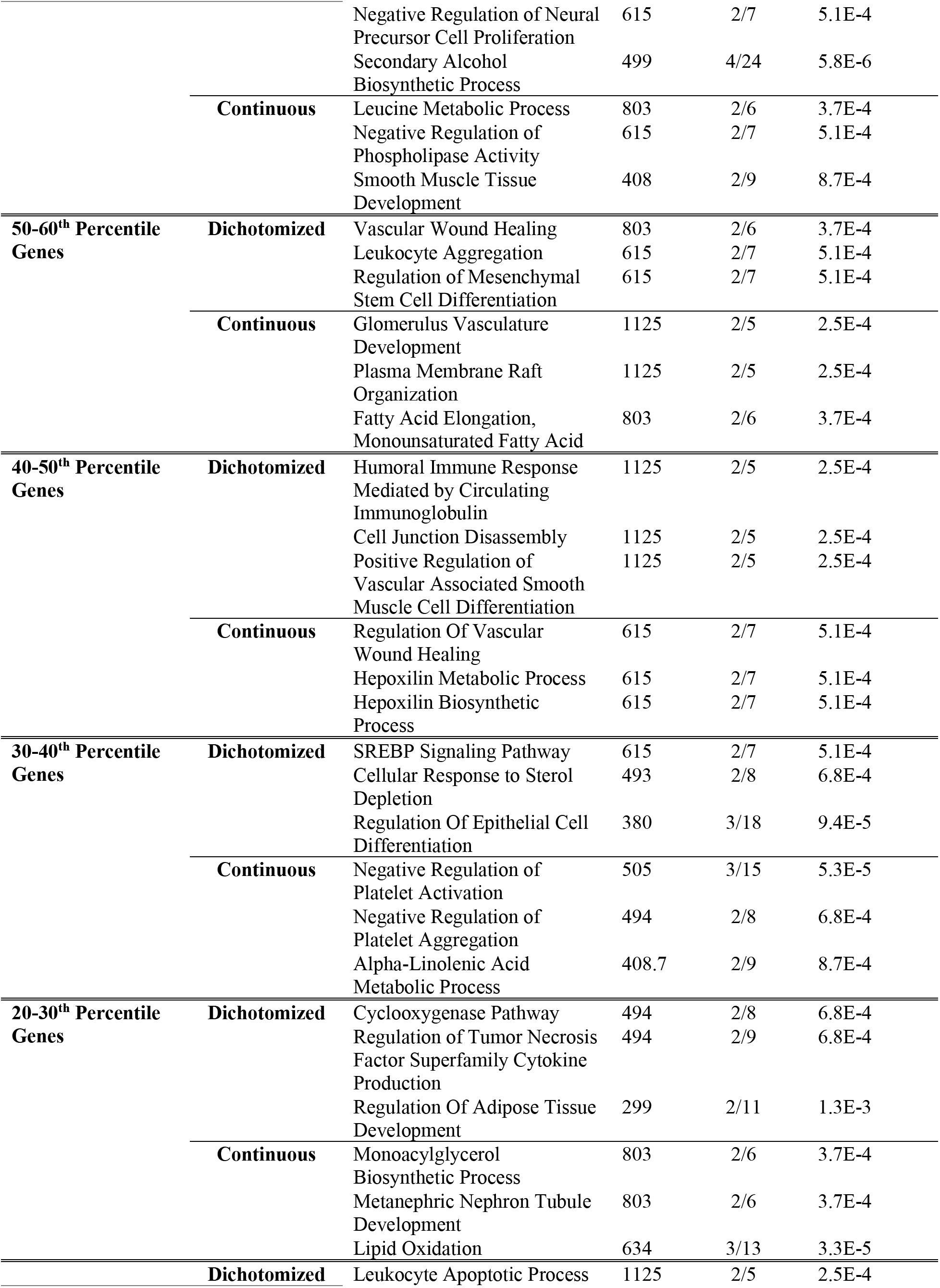
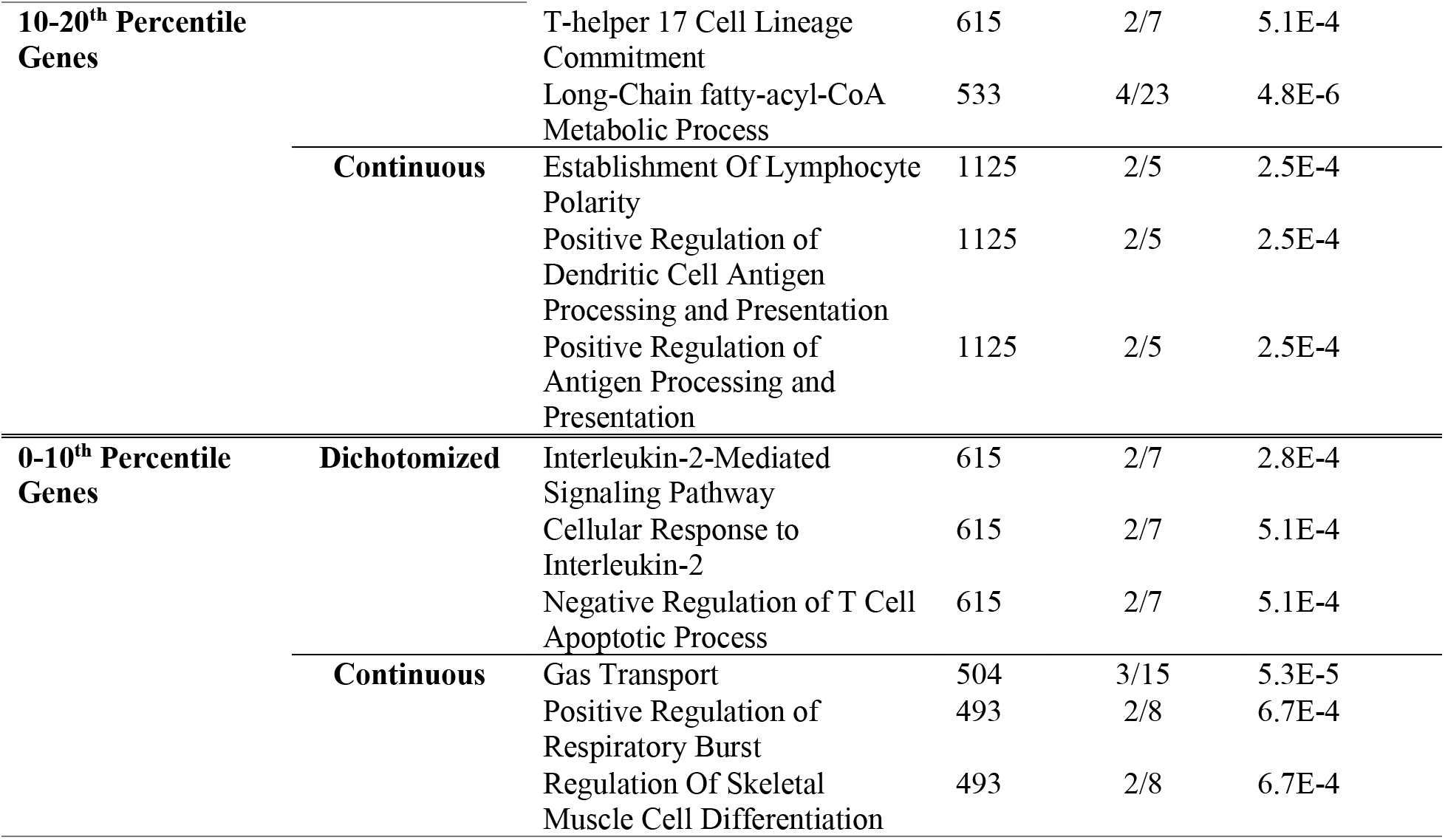
Extended Performance Pathway Analysis. Combined performance statistics for both the dichotomized and continuous models were used to perform a performance stratified pathway analysis. AUC and Spearman coefficient were used to stratify genes in the dichotomized and continuous tasks, respectively. The top 3 pathways, measured via EnrichR using the Go Biological Process 2023 database, are reported for the highest and lowest performance deciles.

**Supplementary Table 3:**
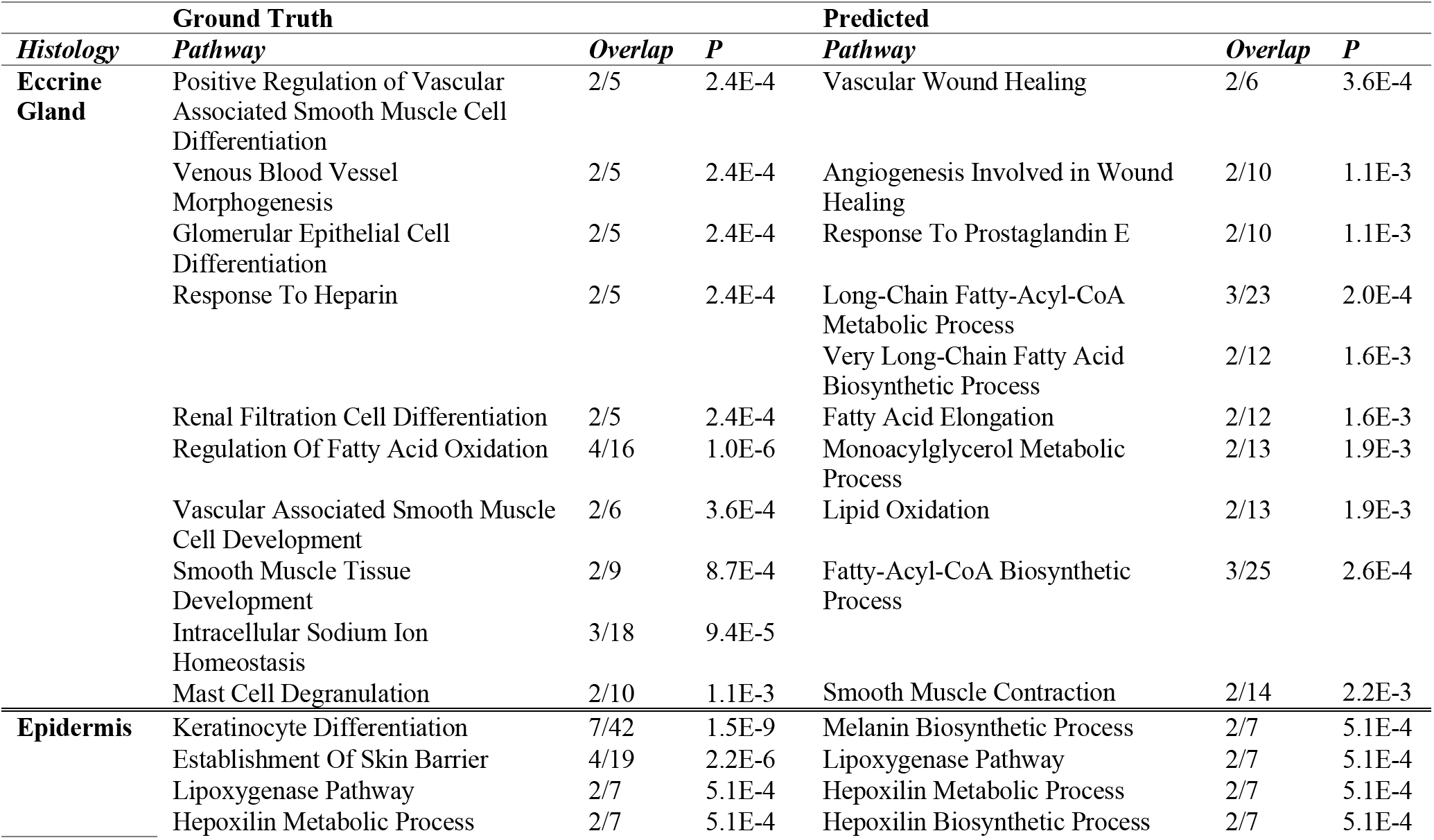
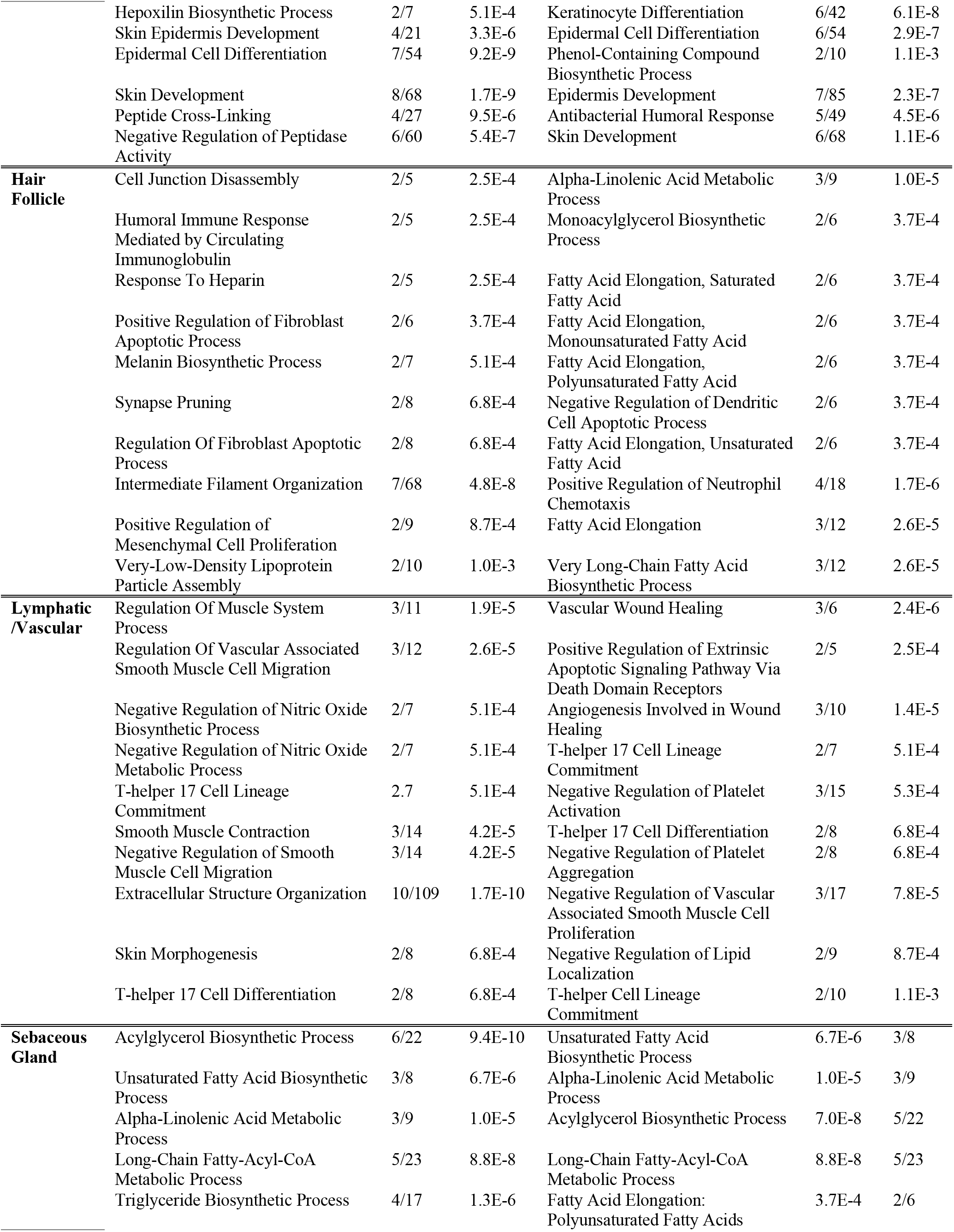
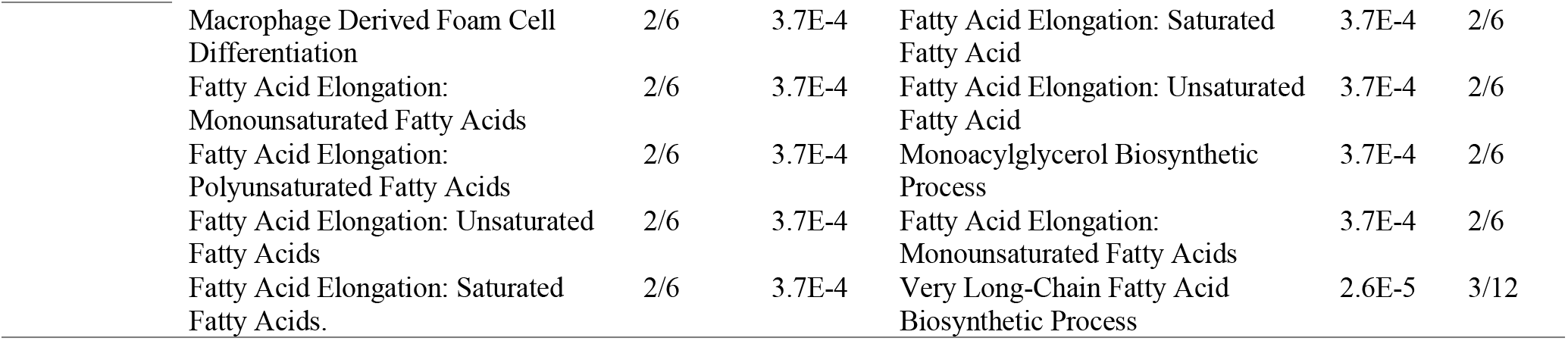
Histological Pathway Analysis for Dichotomized Expression. The top 100 differentially expressed genes for each cluster were established via the Wilcoxon test. These genes were then analyzed using *EnrichR* and the Go Biological Process 2023 database to find the top 10 most salient biological pathways as measured by magnitude and statistical significance (i.e., combined score). These pathways are reported for both the predicted and ground truth data. Bolded pathways are shared between both predicted and ground truth data. Results for sample 14 are displayed for brevity.

**Supplementary Table 4:**
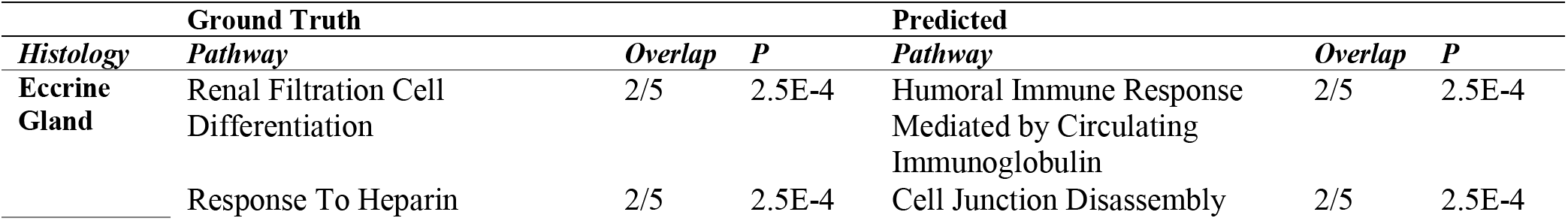
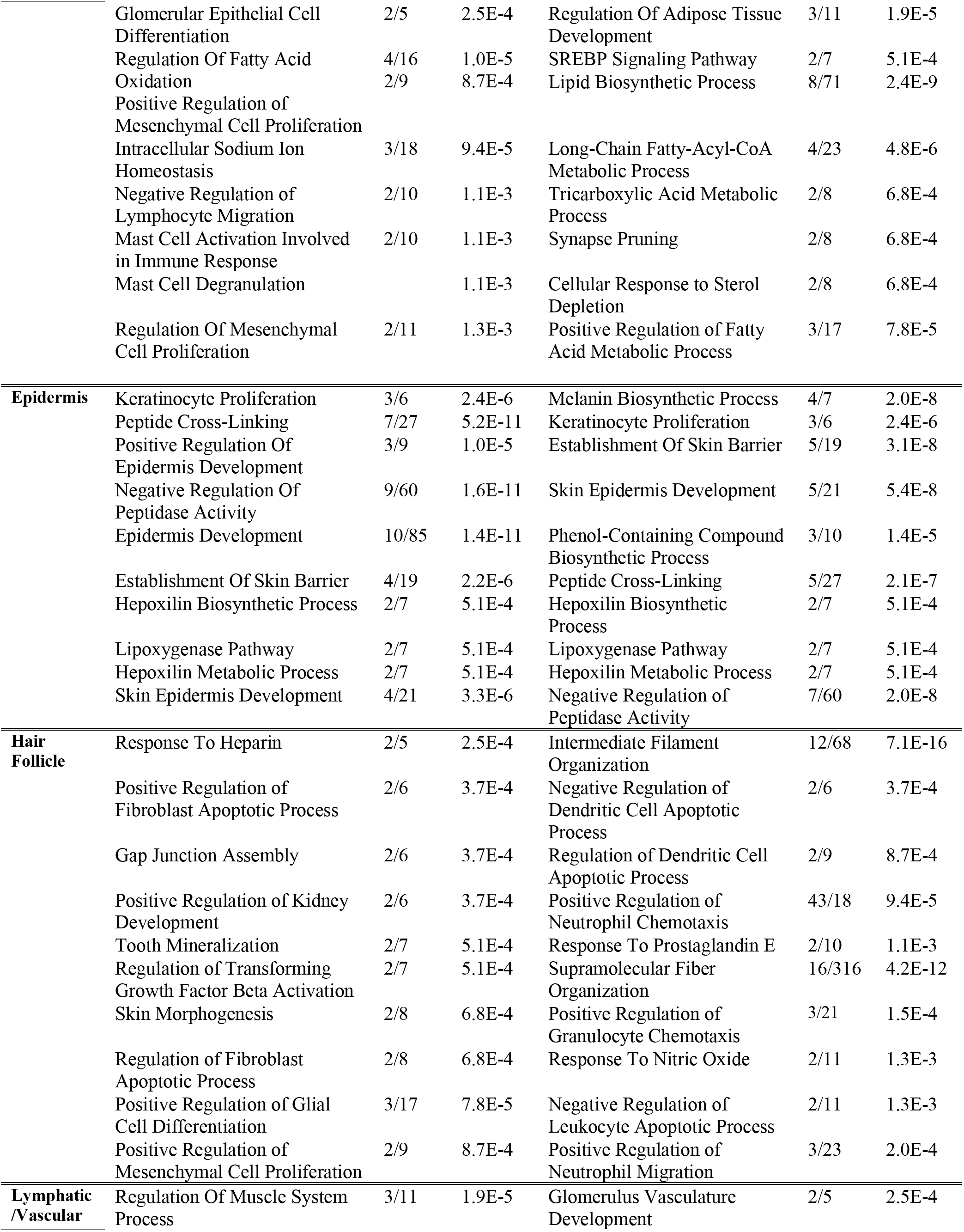
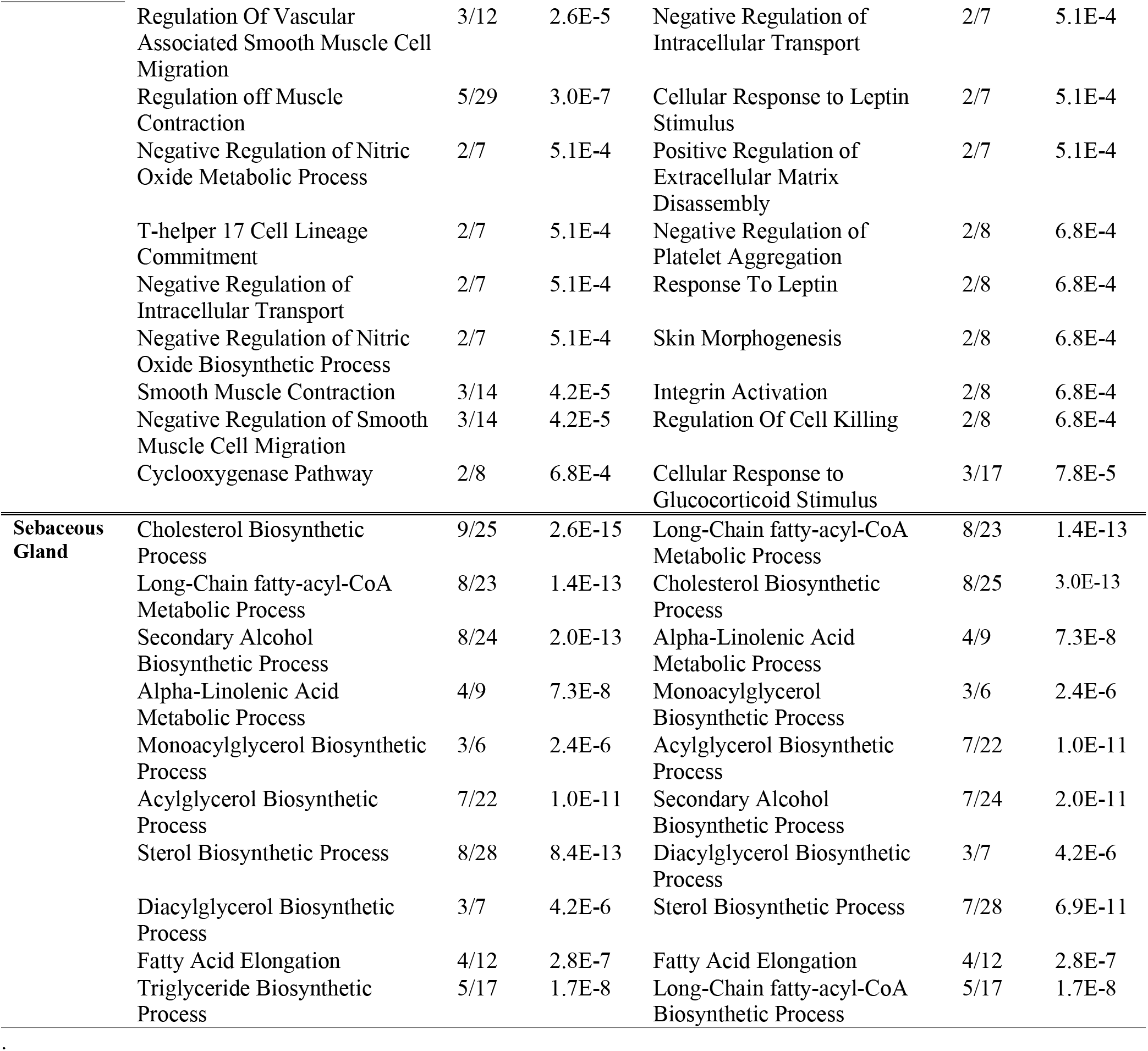
Histological Pathway Analysis for Continuous Expression. The top 100 differentially expressed genes for each cluster were established via the Wilcoxon test. These genes were then analyzed using EnrichR and the Go Biological Pathways database to find the top 10 most salient pathways as measured by combined scores. These pathways are reported for both the predicted and ground truth data. Bolded pathways are shared between both predicted and ground truth data. Results for sample 14 are displayed for brevity.

## Footnotes

† Supplementary materials can be found at the following DOI: https://doi.org/10.5281/zenodo.8197850

